# Correlation of pathogenic factors with antimicrobial resistance of clinical *Proteus mirabilis* strains

**DOI:** 10.1101/2020.02.24.962514

**Authors:** Aneta Filipiak, Magdalena Chrapek, Elżbieta Literacka, Monika Wawszczak, Stanisław Głuszek, Michał Majchrzak, Grzegorz Wróbel, Marek Gniadkowski, Wioletta Adamus-Białek

## Abstract

*Proteus mirabilis* is the third most common etiological factor of the urinary tract infection (UTI). It produces urease, which contributes to the formation of crystalline biofilm, considered to be one of the most important virulence factors of *P. mirabilis* strains, along with their ability to swarm on a solid surface. The aim of this study was to analyze the pathogenic properties of two selected groups of clinical *P. mirabilis* isolates, antimicrobial-susceptible and multidrug-resistant (MDR), collected from hospitals in different regions in Poland. The strains were examined based on virulence gene profiles, urease and hemolysin production, biofilm formation, and swarming properties. Additionally, the strains were differentiated based on the Dienes test and antibiotic susceptibility patterns. It turned out that the MDR strains exhibited kinship more often than susceptible ones. The strains which were able to form stronger biofilm had broader antimicrobial resistance profiles. It was also found that the strongest swarming motility correlated with susceptibility to most antibiotics. The correlations described in this work encourage further investigation of the mechanisms of pathogenicity of *P. mirabilis*.

**Author summary:** *Proteus mirabilis* is widely widespread in environment but also it is responsible for most *Proteus* infections, especially in human urinary tracts. They cause complicated, persistent infections especially due to the ability to form urinary stones. The clinical importance of *P. mirabilis* have been described in the literature many times. However, the role of pathogenic features with correlation to drug resistance require further investigation. In this research we analyzed thee virulence factors in relation to drug resistance of clinical *P. mirabilis* strains isolated from urine. The virulence genes, ureolytic and hemolytic activity, biofilm formation, swarming growth and strains kindship were analyzed. The most important observation was that the strains exhibited a stronger territorialism were kindred to a lower number of other strains, formed weaker biofilm and exhibited a lower resistance to antibiotics. Furthermore, we proved that the strains which were more likely to mutual growth, they were also less similar in the drug resistance profile but exhibited a higher resistance to antibiotics, which can be beneficial for different bacteria living together. We believe that *P. mirabilis* with strong territorialism can represent a wild group of strains with poor experience of antibiotic pressure. The environmental influence (toxins, antibiotics, bacterial neighbors) stimulates the development of a less dispersed community with stronger biofilm, exchange of genes and increase of resistance to antibiotics.

## Introduction

Following *Escherichia coli* and *Klebsiella pneumoniae, Proteus mirabilis* is the third most common etiological factor of the urinary tract infection (UTI) [1–5], being mainly responsible for complicated UTIs or UTIs in long-term catheterized patients. Furthermore, the ability of *P. mirabilis* to form apatite or/and struvite stones in bladder and kidney causes severe pain in patients, and augments therapeutic difficulties [3,6,7]. Besides UTIs, this pathogen develops diverse diseases of the respiratory tract and infections of skin and soft tissue (including postoperative wounds, burns, etc.) [8]. *P. mirabilis* expresses a number of virulence factors, that allow for *e. g*. effective motility against the stream of urine, uptake of nutrients or protection from the host defense system. The most typical are fimbriae, which in general mediate attachment to uroepithelial cells. *P. mirabilis* carries genes of 17 distinct fimbrial structures, the most important being mannose-resistant *Proteus*-like pili (MR/P), *P. mirabilis* P-like pili (PMP), *P. mirabilis* fimbriae (PMF), ambient-temperature fimbriae (ATF) and uroepithelial cell adhesin (UCA) [2,8]. Other significant virulence factors include toxins (HpmAB), iron and zinc uptake systems, proteases and flagella [1,2,4], and urease which hydrolyses urea to ammonia and carbon dioxide. This activity is a substantial source of nitrogen for bacteria and also contributes to the formation of crystalline biofilm that blocks the catheter lumen which is considered one of the most important virulence factors of *P. mirabilis* [3,9–11]. Pathogenicity of *P. mirabilis* strains could also be associated with their ability to swarm on a solid surface [6]. This phenomenon relies on a conversion of short swimmer cells into long, hyper-flagellated swarmer cells, and has been used in a simple Dienes test to differentiate *Proteus* spp. isolates. Its principle is based on the occurrence of boundaries between zones of the swarming growth of non-related strains, while those produced by isogenic strains fuse with each other. However, the background of this effect remains unclear [12].

Over the past decades, clinical strains of *P. mirabilis*, just like other *Enterobacterales*, have become increasingly resistant to antimicrobials that is a serious problem for hospitalized patients [13–16]. In some countries strains with extended-spectrum β-lactamases (ESBLs) or AmpC-like cephalosporinases have spread, which apart from resistance to penicillins and cephalosporins (including oxyimino-compounds), display broad resistance to other anti-infectives [13,17–20]. In Poland, CMY-2-type AmpC producers may account for more than 20% of *P. mirabilis* isolates causing nosocomial infections [21]. The aim of this study was to analyze virulence properties against antimicrobial susceptibility profiles in a collection of selected clinical *P. mirabilis* strains.

## Results

### Antimicrobial susceptibility

Antimicrobial susceptibility profiles of the *P. mirabilis* strains are shown in supplementary data (S1 Table). In general the study sample was pre-selected based on β-lactam susceptibility phenotypes and β-lactamase content, including 22 isolates with CMY-2-like AmpCs and one isolate with a CTX-M-1-like ESBL (all co-producing TEM-1/-2 β-lactamases), two isolates with TEM-1-like enzymes only, and 25 β-lactamase-negative isolates. The β-lactam susceptibility patterns, for 15 CMY producers reported previously [18,22], corresponded well to the β-lactamase content of the isolates, briefly with resistance to penicillins and oxyimino-cephalosporins in AmpC or ESBL producers, and to penicillins only in TEM-1 producers. Resistance to aminoglycosides, fluoroquinolones, co-trimoxazole and/or chloramphenicol occurred frequently in the 25 β-lactamase producers, defining each of these as MDR [23], despite the variety of individual susceptibility patterns and levels of resistance to particular compounds. Otherwise all of the 25 β-lactamase negatives were susceptible to all antimicrobials tested. The lower *in vitro* activity of imipenem, characteristic of *Proteus* and related genera (www.eucast.org), was observed among both MDR and susceptible isolates; however, with higher MICs among the former ones.

### Virulence factors

All of the *P. mirabilis* strains were positive for the studied pathogenicity-related genes (*fliL, mrpA, pmfA, ureC, zapA, hpmA, hpmB*, and *uca*). Eighty-five percent of the isolates produced a transparent zone or color change on the blood agar; however, only eight strains (16%) exhibited the typical β-hemolytic activity. The rate of ureolytic activity was measured in a time course, and it was independent on the bacterial growth rate (Fig 1). All of the strains hydrolyzed urea, but the lowest and the highest level of urease activity in the first and the last hour varied in the range of approximately 30%. The highest increase in urease activity was detected in the third hour of incubation in the Christensen broth. A similar observation was done during the incubation of the agar medium where the enlargement of the ureolytic zone was observed at different rates for individual strains (differences between the strains were from 72% to 58% for the 3rd to 6th hour of incubation, respectively). Generally, despite the differences between strains, the ureolytic activity was stable in the time course.

**Fig 1.**
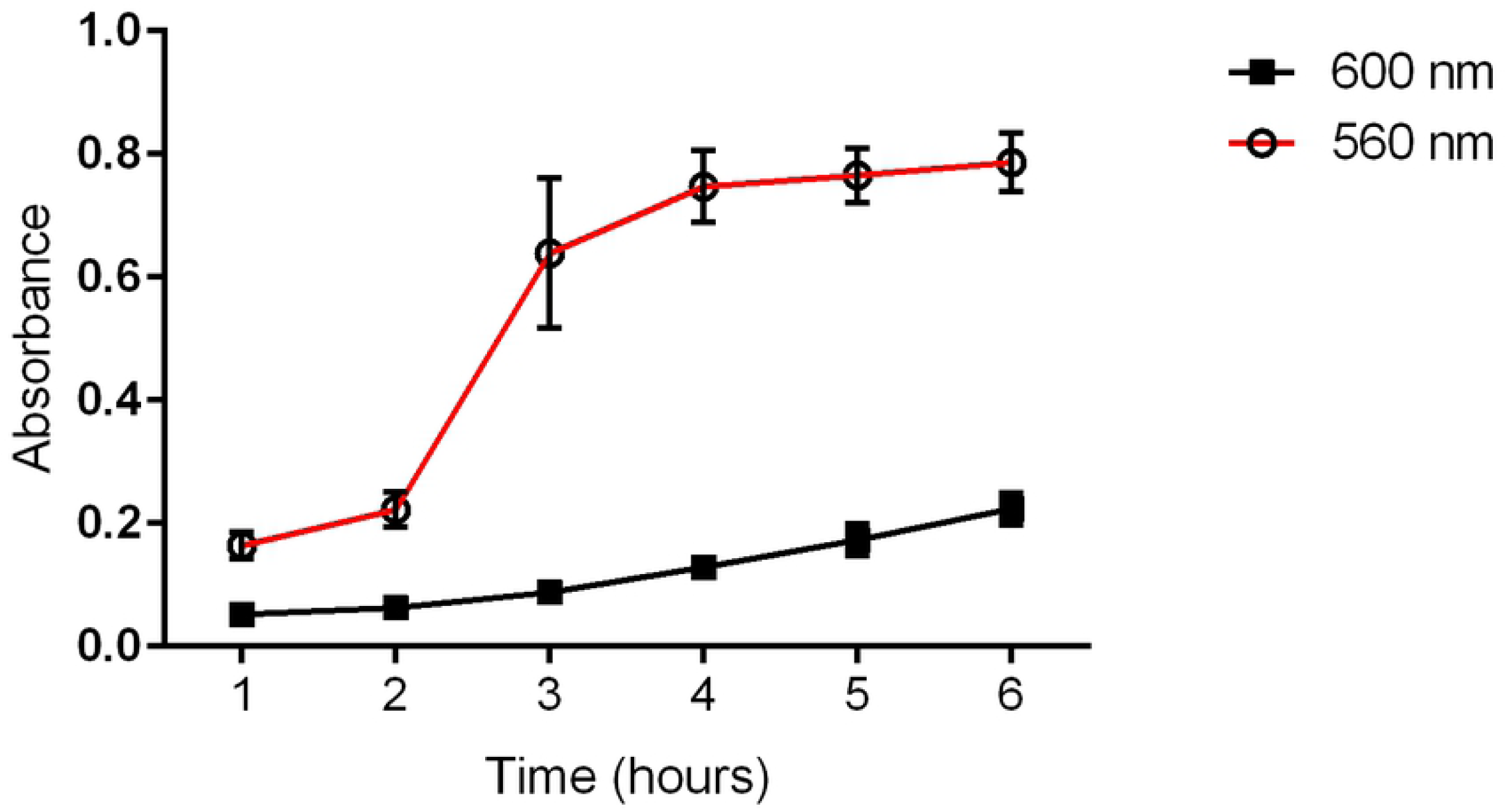
Medium ureolytic activity (560 nm) and bacterial growth (600 nm) of clinical *P. mirabilis* strains. The absorbance was measured spectrophotometrically in each subsequent hour of incubation.

### Swarming motility and the Dienes test

The *P. mirabilis* strains were checked for their swarming motility rate. Twenty percent of the examined isolates exhibited poor swarming growth, 36% showed medium swarming, and 44% displayed intensive swarming. Considering the Dienes test, the strains exhibited kinship with different numbers of other isolates (Table 1). The lowest number of kinship was expressed by four strains that exhibited confluent growth with only three other strains, and the highest number was observed in case of one strain, which was kindred with 31 others.

**Table 1.** Summary of Dienes test results. The differentiation of *P. mirabilis strains* was prepared based on the number of strains which are kindred with others strains.

|  |  |  |  |  |  |  |  |  |  |  |  |  |  |  |  |  |  |  |
| --- | --- | --- | --- | --- | --- | --- | --- | --- | --- | --- | --- | --- | --- | --- | --- | --- | --- | --- |
| **number of strains** | **4** | **4** | **3** | **4** | **1** | **8** | **5** | **7** | **2** | **1** | **1** | **1** | **1** | **2** | **2** | **1** | **2** | **1** |
| **number of kin strains** | 3 | 4 | 5 | 6 | 7 | 8 | 9 | 10 | 11 | 12 | 13 | 14 | 15 | 16 | 20 | 27 | 29 | 31 |

### Biofilm formation

The *P. mirabilis* strains were tested for their ability to form biofilm. The absorbance values of the crystal violet adsorbed on the polyurethane surface were different for individual strains, and these ranged between 0.079 and 1, with the medium absorbance level at 0.4. Only in case of two strains the absorbance level was consistent with the control value. The biofilm formed on the glass positively correlated with biofilm on the polyurethane: the strains unable to form biofilm on the glass exhibited significantly weaker biofilm on the polyurethane, compared to the strains forming stronger biofilm (microcolonies or entire coverslip covered) (Fig 2). Significantly more strains were classified as unable to form biofilm on the glass (42%) than on the polyurethane surface (4%).

**Fig 2.**
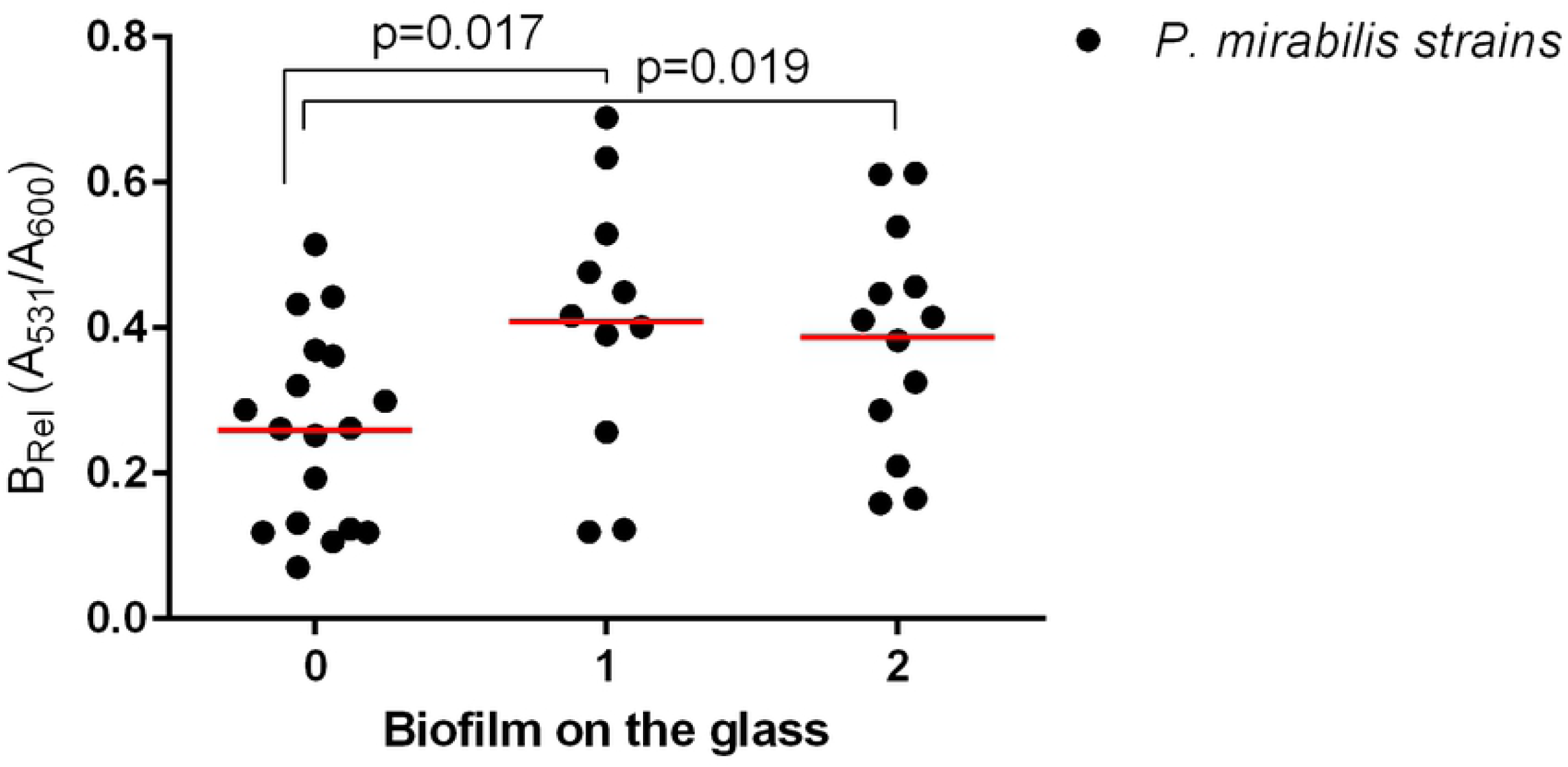
Correlation between biofilms formed on the polyurethane and glass surfaces of clinical *P. mirabilis* strains. Biofilm formed on the polyurethane was measured spectrophotometrically with crystal violet absorbance at 531 nm. Relative biofilm (B_Rel_) was calculated based on the ratio between A_531_ and optical density of the planktonic growth measured at 600 nm to estimate the biofilm formation independent on the bacterial growth rate. Biofilm on the glass surface: lack of biofilm (0), microcolonies (1) and biofilm covering the entire field of microscopic observation (3). The medium values of biofilm were marked with red lines. The unpaired two-tailed T-test was used for a statistically significant difference (p<0.05) (Graph Pad Prism v. 6).

### Correlations

A more detailed comparison of the results obtained in the above-mentioned experiments allowed to observe several dependencies. The kinship of the examined strains correlated with their susceptibility profiles (Table 2). For each strain, the number of kindred strains and the number of not kindred strains were noted, and susceptibility profiles in these groups were analyzed, determining the percentage of the profiles similarity (“A profile”). The “A profile” means the % of susceptibility pattern similarity between a given strain and other kindred or not kindred strains. Forty-four percent of the strains (n=22) revealed significant correlation between kinship and susceptibility profiles. Eighty-six percent of these (19/22) had more similar susceptibility patterns with the group of not kindred strains. The similarity of susceptibility profiles was not observed in the group of kindred strains. Only in three cases the kin strains had similar susceptibility profiles.

**Table 2.** Correlation between kindship of *P. mirabilis* strains measured by Dienes test and median percentage similarity of their antibiotic sensitivity profiles (A profile). The statistically significant (p<0.05) differences in A profiles were assessed with the use of the Mann-Whitney test.

| *P. mirabilis*<br>strain | other strains |  |  |  |  |
| --- | --- | --- | --- | --- | --- |
|  | KINDRED |  | NOT KINDRED |  | p-value |
|  | No. | A profile (%) | No. | A profile (%) |  |
| 368 | 8 | 35 | 40 | 95 | 0,0386 |
| 845 | 9 | 80 | 39 | 50 | 0,0016 |
| 2014 | 9 | 85 | 39 | 40 | 0,0003 |
| 2198 | 14 | 50 | 34 | 70 | 0 |
| 3010 | 28 | 30 | 10 | 57.5 | 0,0174 |
| 5112 | 10 | 37.5 | 38 | 95 | 0,0119 |
| 5618 | 7 | 30 | 41 | 95 | 0,0092 |
| 5653 | 9 | 35 | 39 | 95 | 0,0219 |
| 5663 | 12 | 30 | 36 | 95 | 0,0076 |
| 5777 | 6 | 37.5 | 42 | 95 | 0,0528 |
| 5778 | 7 | 30 | 41 | 95 | 0,0105 |
| 6103 | 15 | 35 | 25 | 55 | 0,0066 |
| 6187 | 6 | 85 | 42 | 30 | 0,0099 |
| 6365 | 8 | 32.5 | 40 | 95 | 0,0194 |
| 6405 | 7 | 35 | 41 | 95 | 0,039 |
| 6521 | 13 | 30 | 35 | 95 | 0,0086 |
| 6771 | 9 | 30 | 39 | 95 | 0,0138 |
| 7101 | 9 | 30 | 39 | 95 | 0,0348 |
| 7104 | 8 | 32.5 | 40 | 95 | 0,0473 |
| 7498 | 7 | 30 | 41 | 95 | 0,016 |
| 7499 | 5 | 30 | 43 | 95 | 0,0163 |
| 7882 | 9 | 35 | 39 | 95 | 0,0004 |

This correlation was more clearly noticed in the analysis of the two groups of susceptible and MDR isolates – (Fig 3). It turned out that kinship with a higher number of strains was observed among MDR strains (p=0.0148). The kinship also correlated with the swarming motility of the *P. mirabilis* strains. The strains with weaker swarming growth were kindred with a higher number of strains compared to those with stronger swarming growth (Fig 4).

**Fig 3.**
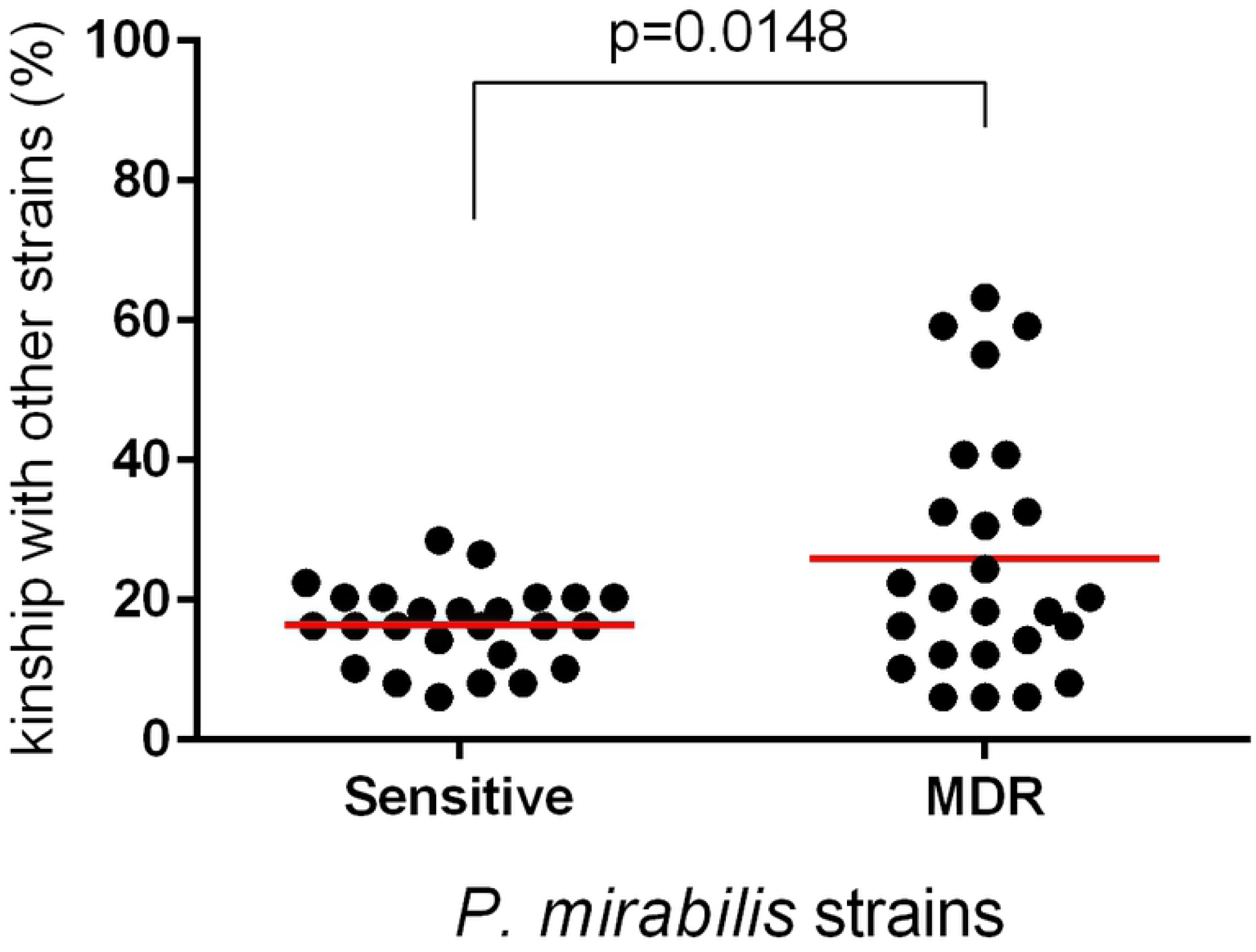
Kinship among sensitive and MDR *P. mirabilis* strains. The medium value was marked with red line. The unpaired two-tailed T-test was used for a statistically significant difference (p<0.05) (Graph Pad Prism v. 6).

**Fig 4.**
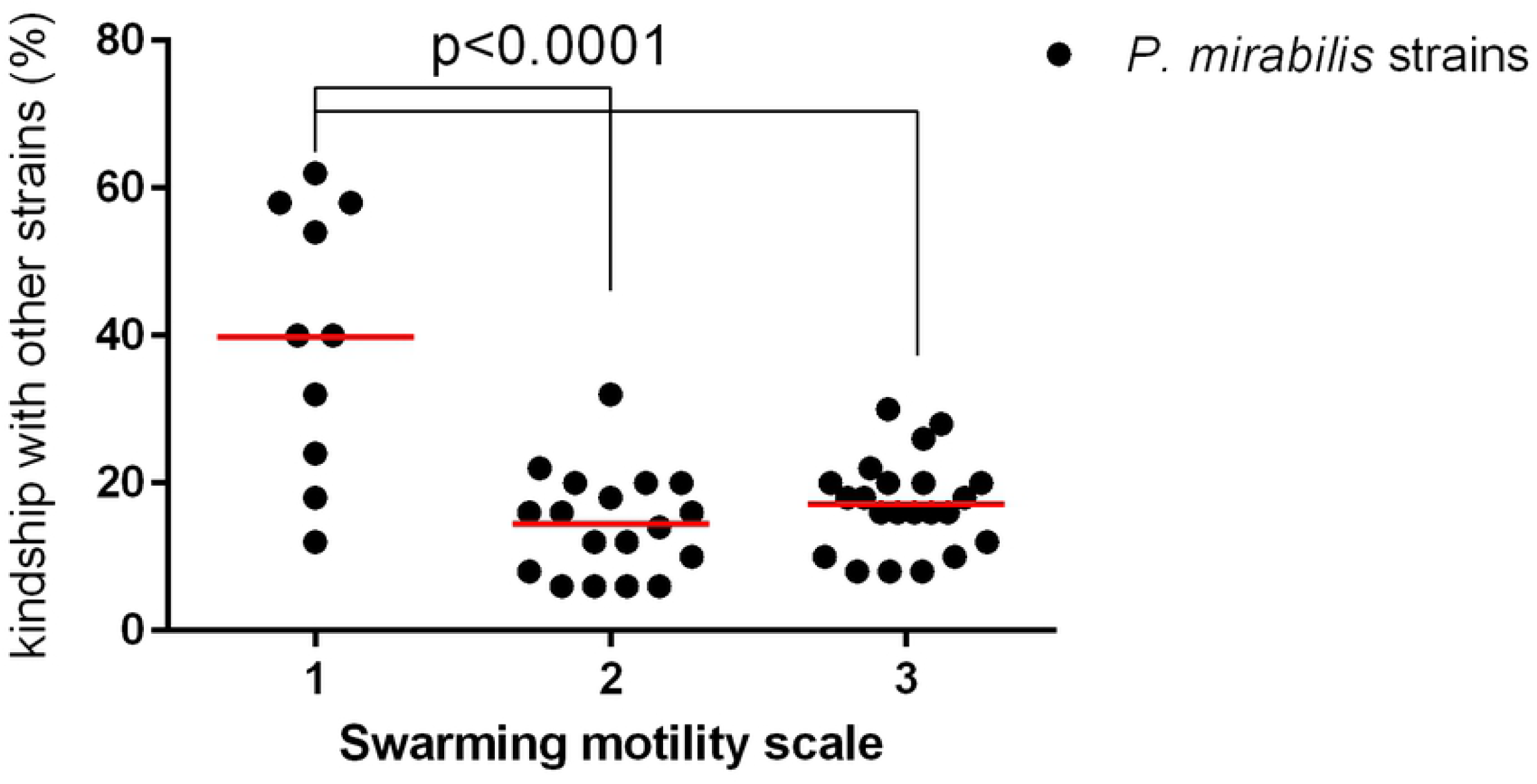
Correlation between kinship and swarming motility among *P. mirabilis* strains. Bacterial strains (n=50) were differentiated based on different swarming motility (in scale of 1, 2 3) and analyzed based on the kinship with other strains (% of examined strains). The medium value was marked with red line. The unpaired two-tailed T-test was used for a statistically significant difference (p<0.05) (Graph Pad Prism v. 6).

The MDR and susceptible groups of strains were compared based on expansiveness of swarming and biofilm strength (Fig 5). Eighty-four percent of susceptible strains exhibited the highest swarming motility rate and these did not include any isolate of the weakest swarming growth. In contrast, 68% of MDR strains exhibited the weakest swarming and only 8% of them showed the strongest motility rate. Additionally, 56% of susceptible strains formed the weakest biofilm, whereas 50% of MDR strains formed the strongest biofilm. Otherwise, the strongest biofilm was formed by only 20% of susceptible strains and the weakest biofilm was found in only 22% of the MDR group. Similar observations were made in the case of the biofilm formation on the polyurethane: the susceptible isolates revealed significantly weaker biofilm when compared to the MDR strains (unpaired Mann-Whitney test, p=0.0371).

**Fig 5.**
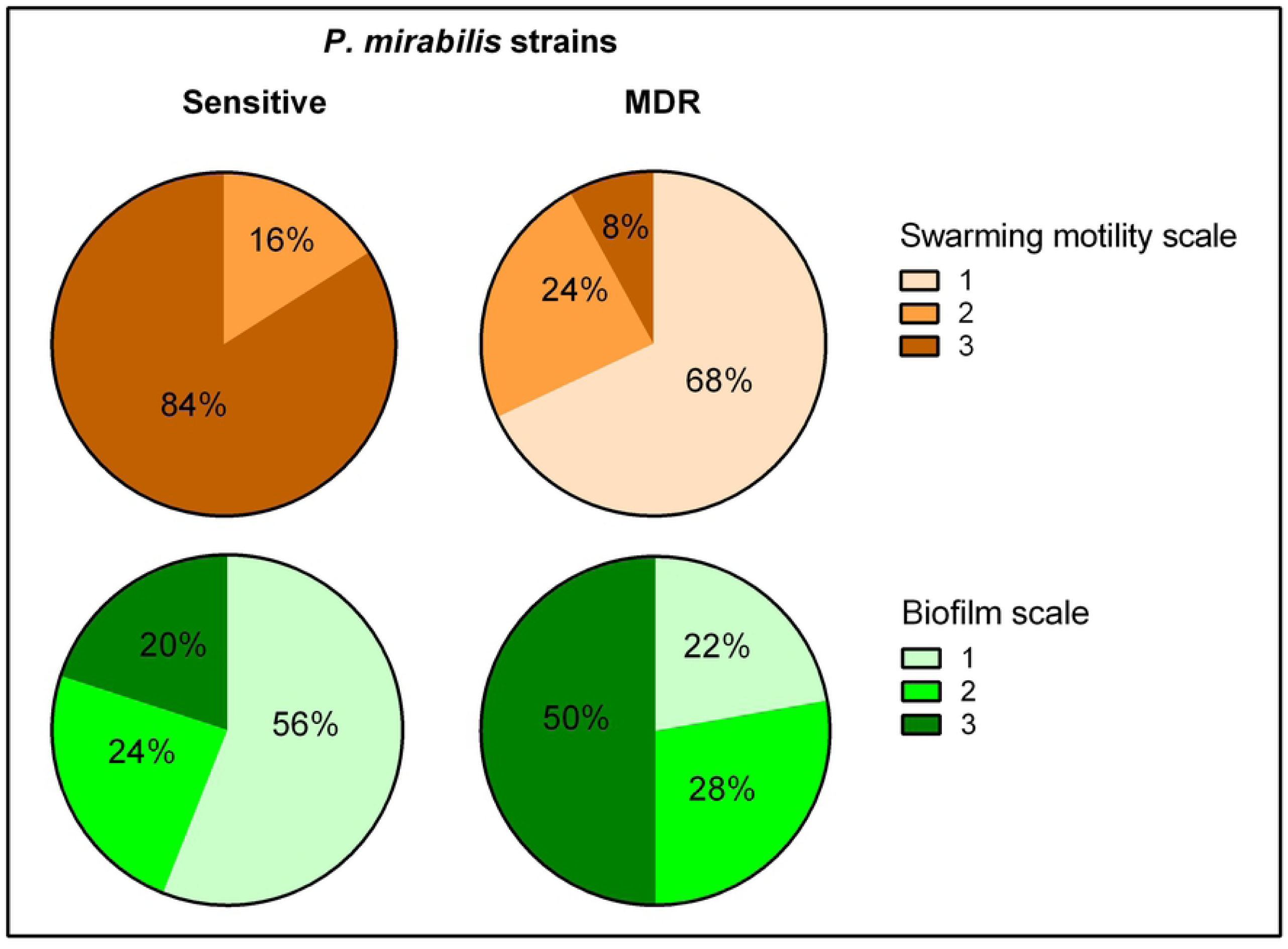
Biofilm and swarming motility among sensitive and MDR *P. mirabilis* strains. The statistically significant difference was observed between sensitive and MDR *P, mirabilis* strains in terms of biofilm formation on the glass (Chi-square test, p=0.0161) and swarming motility (Chi-square test, p=0.0001) measured in three-stage scale (Graph Pad Prism v. 6).

### Differentiation of the isolates

The *P. mirabilis* isolates were differentiated using the Euclidean distance between the MICs. As expected, two distinct clusters were observed in the dendrogram (Fig 6). Cluster 1 contained a highly homogeneous group of susceptible strains, whereas cluster 2 with MDR strains exhibited significantly larger differences in MICs (two tailed T-test, unpaired, p<0.0001). These clustering with clear distinction of bacterial antibiotic sensitivity level confirmed the correlations described above but here we proved their interdependence. Cluster 1 represents the strains kindred with fewer strains compared to cluster 2 (two tailed T-test, unpaired, p=0.017), i.e. the cluster 2 strains exhibited kinship to each other more often than those in cluster 1. Moreover, cluster 1 included strains forming weaker biofilm, both on the polyurethane (two tailed T-test, unpaired, p=0.055) and on the glass (two tailed T-test, unpaired, p=0.015), and exhibiting statistically significant more expansive swarming motility (two tailed T-test, unpaired, p<0.001) compared to cluster 2. There was no difference either in the level or rate of urease and hemolysin production between the two clusters.

**Fig 6.**
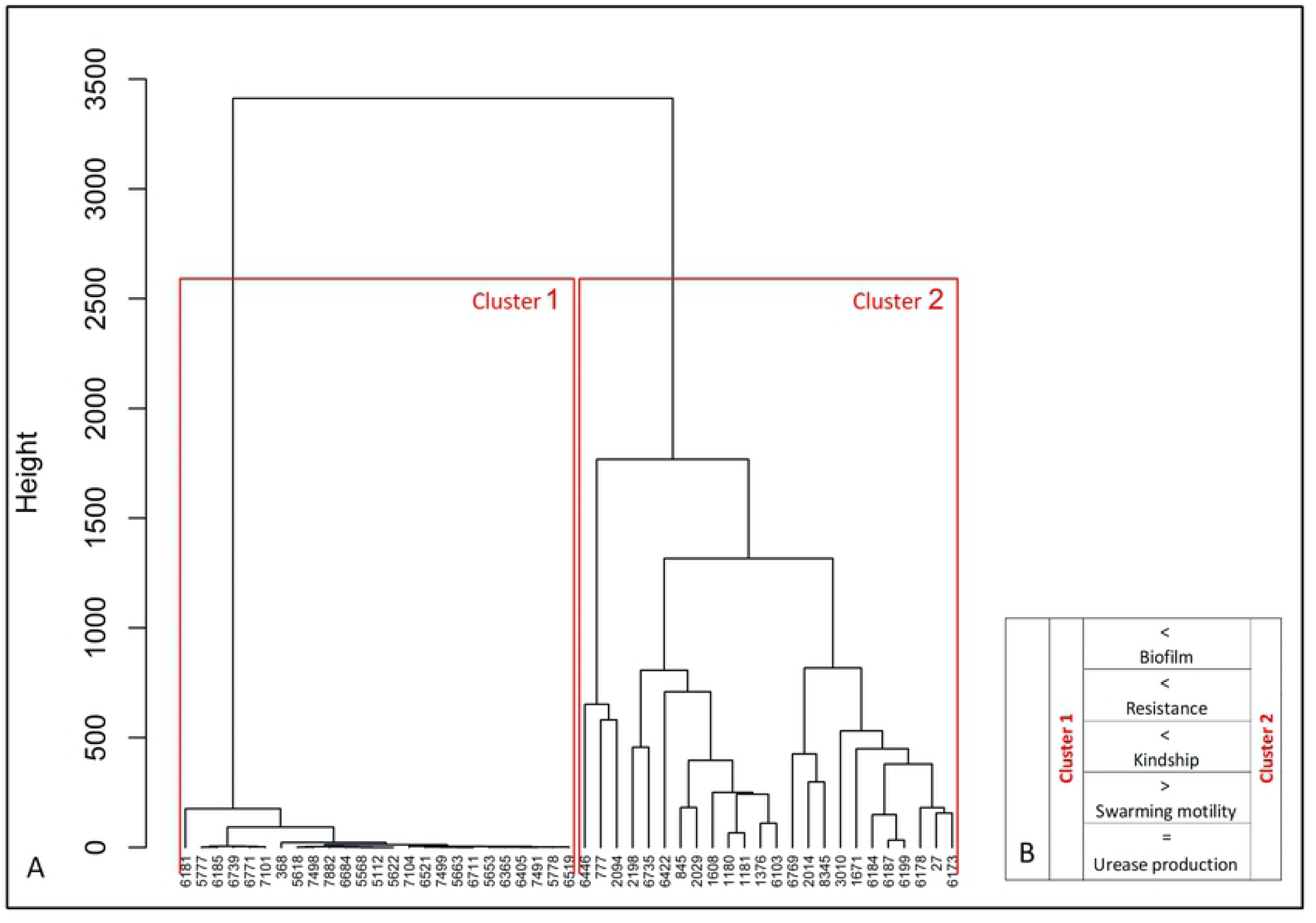
Differentiation of *P. mirabilis* strains based on the profiles of MICs of antibiotics. The similarities of MICs of antibiotics were used for the dendrogram (A) developed by Ward’s agglomeration. The schematic characteristic of cluster 1 and cluster 2 was added (B) based on the bacterial features, where >, < higher/stronger,= the same level.

## Discussion

The study was focused primarily on pathogenicity factors of *P. mirabilis* isolates, recovered from hospital UTIs in different regions in Poland, and selected based on their susceptibility profiles. Previous reports have documented a clear epidemiological success of the MDR AmpC-producing strains in Greece, Italy and Poland, where they accounted for approximately 20% of nosocomial clinical *P. mirabilis* isolates [18,22]. This prompted us to check for other factors, including virulence properties, that might have contributed to their effective dissemination in hospital environments.

Owing to specific virulence factors, *P. mirabilis* is particularly troublesome for catheterized patients and is responsible for complicated UTI with the development of urinary stones [24–26]. In our study we analyzed the presence of virulence factor genes, starting with *fliL*, representative for the class II flagellar operon (*fliLMNOPQR*) [27], which is involved in the swarming motility. Other genes included those determining the mannose-resistant *Proteus*-like adhesion (*mrpA*), *P. mirabilis* fimbriae (*pmfA*) and uroepithelial cell adhesion (*uca*). They play a crucial role in the catheter-associated biofilm formation, and the bladder and kidney colonization, respectively [10,26,28]. The remaining tested genes *zapA, hpmA, hpmB* and *ureC* are engaged in the immune system evasion and/or iron acquisition [24], with *zapA* also involved in the swarmer cell differentiation and swarming behavior [29]. All of the examined *P. mirabilis* strains carried all of the virulence genes addressed, which is consistent with previous reports [1,30]. Nevertheless, they underline the often observed poor expression and downregulation of virulence genes. The *P. mirabilis* chromosome is strongly conservative and the location of the analyzed genes seems to be more stable [1] compared to *E. coli* [31].

Despite the common presence of *fliL* and *zapA*, we observed a poor or even lack of the typical swarming growth in approximately 20% of the isolates that might be due to their low expression and/or correspond to the significant role of other genes in this type of motility [32–36]. This mechanism is also associated with phylogenetic relationships among *P. mirabilis* strains [29,37]. We noticed that the strains with more expansive swarming were related to less of the other strains in the Dienes test. The demarcation line is still rather poorly understood, and probably regulated by multiple mechanisms, like the type VI secretion system, operons *ids*ABCDEF and *idr*ABCDE, and the hpc-vgrG effector [38–40]. However, factors responsible for the disability of strains to extensively swarm in place of the mutual cohabitation with other strains have not been unambiguously identified. Budding et al. [41] suggested that this characteristic might reflect environmental competition between *P. mirabilis* strains which might also have implications in hospital settings.

No correlation between the urease and hemolysin expression levels and the other pathogenicity properties or susceptibility profiles was observed. The ureolytic activity peaked quickly during culture incubation and was independent on the type and rate of the bacterial growth, *i*.*e*. in broth and on agar, being so similar in both swimmer and swarmer cells. This activity is critical in the formation of bacteria-induced stones, particularly dangerous for long-term catheterized patients [1,42]. Urease is also involved in forming crystalline biofilms on the catheter [24]; however, we did not observe any correlation between the enhanced urease expression and stronger biofilm.

The ability to form biofilm varies remarkably, even among strains of the same species [43–45]. Our results (Fig 7) indicate that the *P. mirabilis* biofilm may be stronger than that of *E. coli* [46]. That may mean that *P. mirabilis* grows faster, providing greater biofilm yield, which might be important during the host invasion. The decrease of the *P. mirabilis* growth rate might inhibit the biofilm formation, unlike in *E. coli*. We also observed that the biofilm formation on the glass correlated proportionally with that on the polyurethane, which was again in contrast to *E. coli* [46]. Czerwonka et al. [11] reported that high hydrophobicity of the *P. mirabilis* cell surface correlated with low biofilm amount, which is important for hydrophobic surfaces like glass. It was proved that hydrophilic catheters may prevent catheter-associated UTIs [47], so it would be advisable to evaluate hydrophobicity rates that are characteristic for the majority of *P. mirabilis* clinical strains.

**Fig 7.**
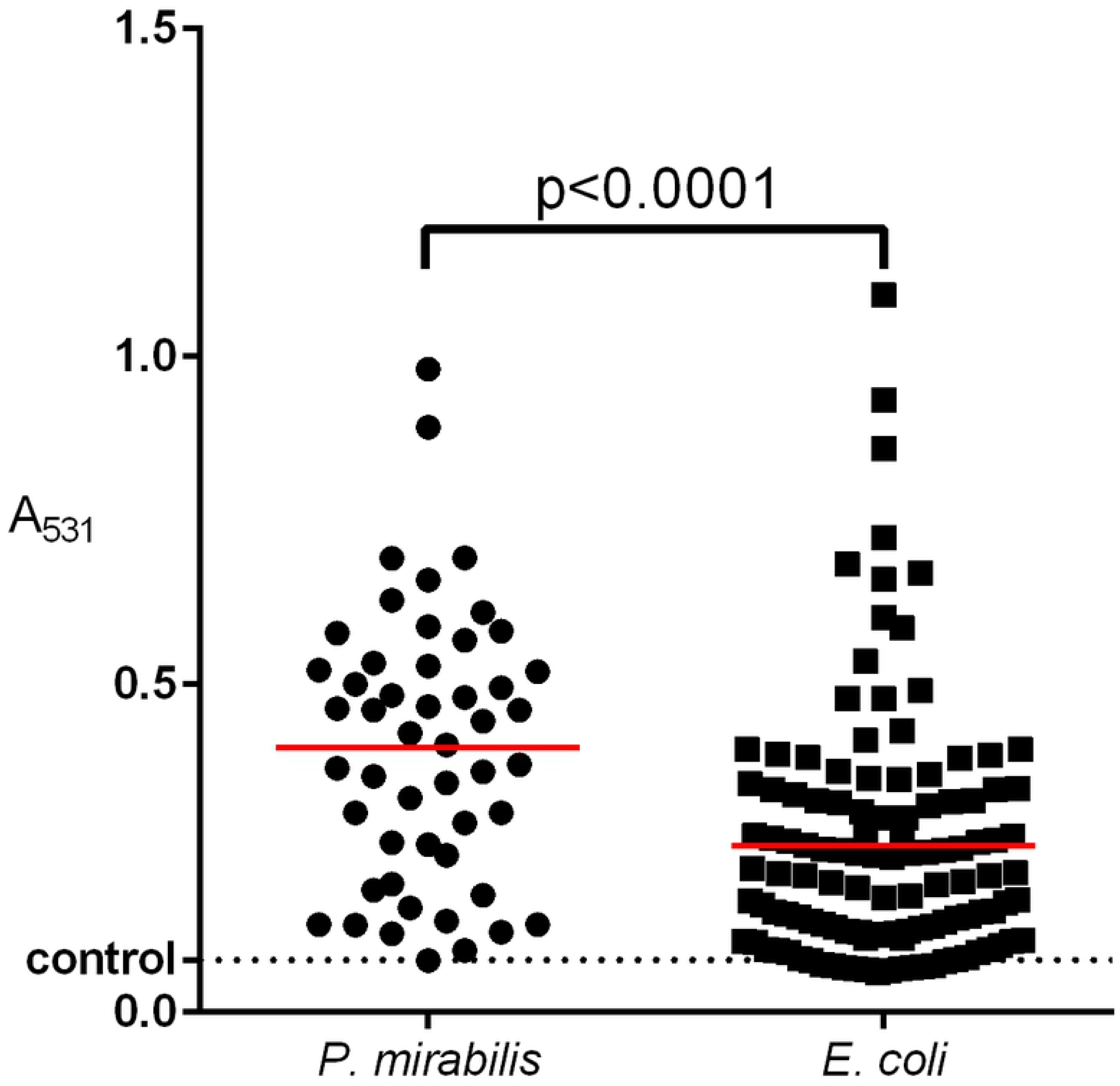
Comparison of biofilm formation between clinical *E. coli* and *P. mirabilis* strains isolated from urine of hospitalized patients. The biofilm was measured in triplicate by crystal violet absorbance at 531 nm. The unpaired two-tailed T-test was used for a statistically significant difference (p<0.05) (Graph Pad Prism v. 6).

The statistical analysis allowed us to observe specific correlations between virulence-associated properties and antimicrobial susceptibility profiles. We found that stronger swarming was significant among susceptible strains and *vice versa*. It was not confirmed in case of tazobactam, the effect of which is independent of the swarming motility [48]. Similar tests were conducted by Auer et al. [49] who tested swarmer cells for sensitivity to wall-modifying antibiotics. They found that thickness of the peptidoglycan makes swarmer *P. mirabilis* cells more sensitive to the osmotic pressure and thus the cell wall-targeting antibiotics. Considering that swarming is an intrinsic feature of *P. mirabilis*, the acquisition of certain antibiotic resistance mechanisms by its individual strains may affect their growth properties, including the ability to transform from swimmer to swarmer cells. Tipping and Gibbs [50] have recently provided a new evidence about the bacterial behavior under the pressure of Ids system which mediates in territorial exclusion among *P. mirabilis* strains. IdsE and IdsD proteins are signaling molecules which are recognize each other themselves among kindred strains. Disruption of this phenomenon leads to creation of a demarcation line and inhibition of the mutual growth of neighbored strains or decreased expansion of one of them with pronounced swarming growth of more aggressive strain. They also observed that mutants without the expression of *IdsE* exhibited an increased tolerance to ampicillin and kanamycin. This recognition signals also need flagellar regulation. Their conclusions are consistent with our observation that strains with weaker swarming growth were more often kindred with other strains and had decreased tolerance to antibiotics, and *vice versa*. The presented observation and other reports encourage further research into this still poorly explored phenomenon.

Susceptibility to antibiotics also correlated with biofilm formation level. In general, we observed that MDR strains formed stronger biofilm, both on the polyurethane and on the glass, compared to susceptible isolates. It has been well-evidenced that biofilm increases resistance to antimicrobials, mainly due to extracellular matrix and weaker penetration of the compounds into the bacterial community [51,52]. However, this is not the only one possible mechanism because some antibiotics penetrate efficiently through biofilm [53]. It has been suggested that some components of the biofilm matrix may affect the activity of antibiotics, like *e*.*g*. exopolysaccharide Psl of *P. aeruginosa*, which inhibits contacts with bacterial cells by electrostatic interactions [54]. The accumulation of bacterial cells can amplify the activity of β-lactamases and other resistance mechanisms, and the downturn of the cellular metabolism inside the biofilm may contribute to resistance to antibiotics [55]. Yet another explanation of the biofilm’s contribution to resistance is the persister phenomenon. Persisters have been described as dormant cells arising within bacterial biofilms with high tolerance to antibiotics [56]. Planktonic persisters are cleared by the host immune response, while biofilm persisters are shielded from host defense [57] and may cause a relapse of infection. Previously we observed that amoxicillin increased the biofilm formation in *E. coli* [31], possibly due to cell wall modifications and induced bacterial stress. It should be underlined that in this study we did not experimentally address changes of antibiotic tolerance during biofilm formation in real time. We only observed that MDR strains formed stronger biofilm, which may have resulted from the presence and/or expression of non-defined factors.

The Ward’s agglomeration test applied for the analyzed MICs allowed for deeper differentiation of the strains. As expected, the isolates were separated into two clusters: cluster 1, grouping the susceptible isolates, and cluster 2, comprising the MDR ones (Fig 6). Moreover, the correlations described above were clearly demonstrated in these two groups. The antibiotic resistance correlated with pathogenic properties of *P. mirabilis* differently than in case of *E. coli* [58]. A similar study of pathogenic properties of *P. mirabilis* was reported by Stańkowska et al. [59] who differentiated the swarming growth rate, and ureotylic, proteolytic and hemolytic activities, calculating the relative virulence index based on cumulative scores for these activities. However, our study showed that virulence properties may be expressed alternatively by *P. mirabilis* strains. Biofilm and swarming growth seemed to be antagonistic to each other, ureolytic activity was similar in all of the isolates, and hemolysin was rarely detected, suggesting lower relevance of the relative virulence index.

To sum up, we excluded the potential rule, that kindred strains which not form a demarcation line will have a similar drug resistance profile. The further discussion indicates the need to rethink the applied concept of kinship in relation to the demarcation line, especially that Tipping and Gibbs [50] emphasize the complex regulation of gene expression involved in this phenomenon. We showed that kindred, or rather compatible strains were less similar in terms of individual detailed antibiotic profiles, expressing also broader resistance and stronger biofilm formation, but less ability to swarm. This compatibility with a larger number of strains might promote mixed colonization or infection, and thus increase the horizontal gene transfer, producing better-adapted organisms that are harder to eradicate [54,55]. Otherwise, the strains, which exhibited frequent incompatibility with others – they revealed also broad susceptibility to antibiotics and stronger swarming motility. These might represent a wild-type group of strains of low or no pre-exposure to antimicrobials. β-Lactams, belonging to the 1^st^ line drugs for the UTI treatment, might select resistant organisms, including the MDR strains with resistance also to other antimicrobial classes. That would explain their clustering in dendrogram. Antibiotics can stimulate the mechanisms of adaptation to unfavorable condition. In the previous study we observed how the antibiotics changed the genetic virulence profiles of uropathogenic *E. coli* strains [31]. In case of *P. mirabilis* a similar phenomenon is also possible but in different features like communication and compromises between *P. mirabilis* strains. Here, for the first time, we described the specific correlations between the pathogenic properties of *P. mirabilis* strains, which encourage further investigation of swarming growth, territoriality mechanism and other pathogenic properties.

## Materials and Methods

### Bacterial strains and antimicrobial susceptibility testing

Fifty non-duplicate clinical *P. mirabilis* isolates deposited in the National Medicines Institute in Warsaw, Poland, were used in this study. They were recovered from urine of patients treated in 24 hospitals in 18 Polish cities from 1998 to 2004. Apart from the broad geographic distribution, the isolates varied in antimicrobial susceptibility, and included 15 multidrug-resistant (MDR) isolates with CMY-2-like AmpC cephalosporinases (CMY-12, −14, −15, −45) plus TEM-1/-2-like broad-spectrum β-lactamases, reported previously [18,21,22]. The 35 remaining isolates were selected based on their β-lactam susceptibility phenotypes, β-lactamase isoelectric focusing (IEF) patterns and β-lactamase gene PCR profiles, determined as described earlier [21,22]. These comprised eight isolates resistant to oxyimino-cephalosporins, including seven further CMY-2-like AmpC and one CTX-M-1-like ESBL producers (all with TEM-1/-2), two ampicillin-resistant isolates with TEM-1-like enzymes, and 25 isolates fully susceptible to β-lactams with no β-lactamases detected in IEF. Minimal inhibitory concentrations (MICs) of 19 antimicrobials (S1 Table) were evaluated by broth microdilution. The methodology and results interpretation were in accordance with the European Committee on Antimicrobial Susceptibility Testing (EUCAST; www.eucast.org), and in case of cefoxitin, with the Clinical Laboratory Standards Institute [60].

### Virulence genes detection

Bacterial DNA was isolated and purified with the GenEluteTM Bacterial Genomic DNA kit (Sigma-Aldrich, St. Louis, Missouri, USA). PCR was used for the identification of six virulence factor genes (*fliL, mrpA, pmfA, ureC, zapA, hpmA, hpmB, uca*). PCRs were performed in 25-μl reaction mixtures containing 12.5μl of DreamTaq™ Green DNA Polymerase Master Mix (2x) (Thermo Fisher Scientific, Waltham, Massachusetts, USA), bacterial DNA (1ng) and 100pmol of each primer. Sequences of primers, their annealing temperatures and amplicon sizes are shown in Table 3. Cycling conditions were as follows: denaturation at 94°C for 2 min, followed by 30 cycles of 1 min at 94°C, 1 min at varying annealing temperature, and 1 min at 72°C, followed by 5 min at 72°C.

**Tab. 3.**
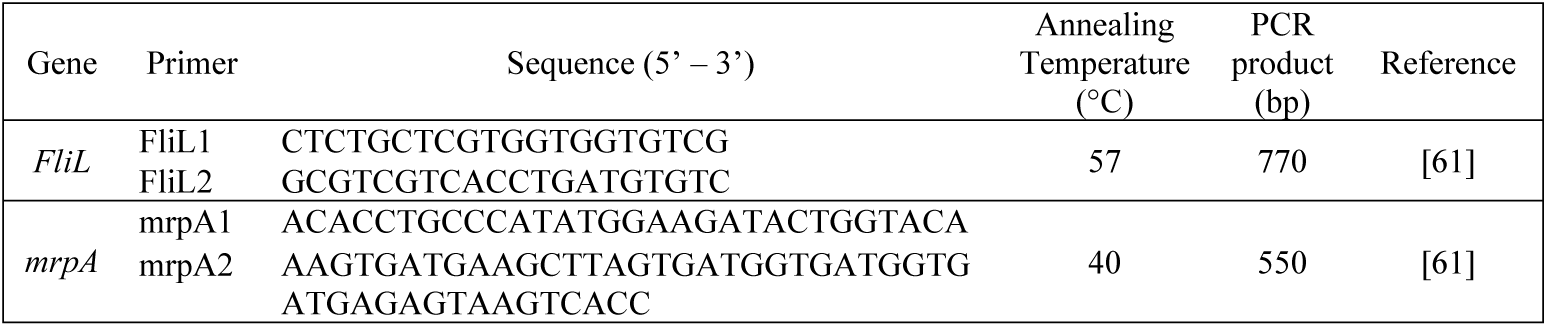
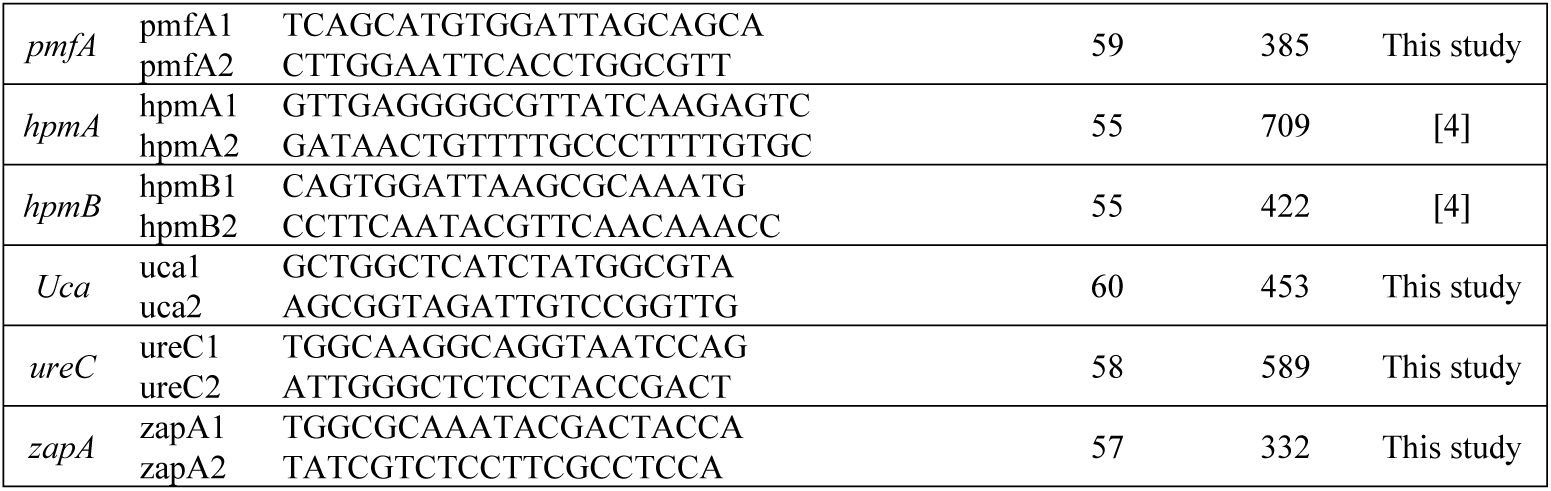
Oligonucleotides used for PCR.

### Hemolysin and urease production

β-Hemolysis properties of *P. mirabilis* strains were determined by observing clear zones around bacterial colonies on blood agar supplemented with 5% (v/v) bovine blood after 24 or 48 hours of incubation in 37°C [62].

The ureolytic activity in the Christensen broth was analyzed according to the method described previously [63] with some modifications. A fresh bacterial inoculum (0.5 McFarland) was diluted 1:100 in the Christensen broth and incubated with shaking (200 rpm) at 37°C for 6 hours. After each hour of incubation, the absorbance of the centrifuged supernatant was measured spectrophotometrically at 560nm. Additionally, the ureolytic activity was observed on the Christensen agar. Briefly, 10μl of a fresh bacterial inoculum (0.5 McFarland) was dropped on the center of an agar plate. The culture was incubated at 37°C for 7 hours and the medium color change (from yellow to pink) was indicative of the urea hydrolysis. The diameter of a pink color zone around the culture was measured after each hour of incubation, starting from the third hour. The dynamics of the ureolytic activity was estimated based on the percentage difference of the activity values between the previous and the next hour of incubation.

### Swarming motility and the Dienes test

The Dienes test was performed according to the protocol described previously [64], with some modifications. Ten microliters of overnight broth cultures were inoculated on Luria Bertani agar plates. Four aliquots of individual cultures were applied onto a single plate at equal distances (about 1cm) from each other, pre-dried for 15 minutes at room temperature, and incubated at 37°C for 24 hours. Bacterial strains showing a clear band in between (Dienes demarcation line) were interpreted as unrelated, while those without the Dienes line were regarded as related to each other. Additionally, the expansiveness / rate of swarming motility was evaluated for each strain by its proportional coverage of the plate after the overnight incubation, compared to other strains cultured on the same plate. The expansiveness was classified into three categories: category 1, weak swarming (coverage <5%); category 2, medium swarming (5% - ≤ 25%); and category 3, intensive swarming (25% - ≤ 50%). The bacterial behavior in each combination of strains was tested in triplicates.

### Biofilm formation

The ability of *P. mirabilis* strains to form biofilm was analyzed on glass and polyurethane surfaces according to the method described previously [46]. The biofilm on the glass was stained with SYTO® 9 solution and propidium iodide, according to the manufacturer’s protocol (Filmtracer™; Invitrogen, Carlsbad, California, USA), and then observed with an epi-fluorescence microscope (Axio Scope.A1; ZEISS, Oberkochen, Germany). Five of the most representative images were photographed. All strains were classified into three groups: group 1, lack of biofilm, single cells observed; group 2, single microcolonies of biofilm; and group 3, biofilm covering the entire coverslip. The biofilm formation on the polyurethane was measured spectrophotometrically at 531nm after overnight incubation of bacterial strains in Luria Bertani broth and crystal violet staining. The strains unable to form biofilm were classified based on the crystal violet adsorption on the polyurethane measured as A_531_<0.08. This value corresponds with the control incubation of crystal violet without bacteria. The relative biofilm (B_Rel_) was estimated as the proportion between biofilm formation level and bacterial growth (A_531_/A_600_).

### Statistical analysis

The group comparisons of *P. mirabilis* strains were carried out using the Fisher exact test for categorical variables, the unpaired t-test for quantitative, normally distributed variables or the Mann-Whitney test for quantitative, non-normally distributed variables (normality of distribution was checked with the Shapiro-Wilk test). The cluster analysis was performed using Euclidean distance and Ward’s method. Statistical tests were two-tailed and p-value of less than 0.05 was considered significant. All statistical analyses were performed using R (version 3.1.2; The R Foundation for Statistical Computing, Vienna, Austria) and Graph Pad Prism v. 6 (San Diego, CA, USA).

## Acknowledgments

We would like to thank Ewa Gwizda for the contribution during the laboratory work of the experiments.

## Supporting information

**S1 Table. Susceptibility of the *P. mirabilis* isolates used in the study**. MICs Minimal Inhibitory Conentrations of antibiotic, - lack of β-lactamases, ^*a*^ – the MICs values shown in bold, italic and normal style correspond to resistance, intermediate resistance and susceptibility, respectively, ^*b*^ – abbreviations: AMC; amoxicillin-clavulanate; AMK, amikacin; AMP, ampicillin; ATM, aztreonam; CAZ, ceftazidime; CHL, chloramphenicol; CIP, ciprofloxacin; CTX, cefotaxime; FEP, cefepime; FOX, cefoxitin; GEN, gentamicin; IPM, imipenem; MEM, meropenem; NET, netilmicin; NOR, norfloxacin; PIP, piperacillin; SXT, trimethoprim-sulfamethoxazole; TOB, tobramycin; TZP, piperacillin-tazobactam.

**S2 Table. The raw data of the research results**.

